# Decoding spontaneous pain from brain cellular calcium signals using deep learning

**DOI:** 10.1101/2020.06.30.179374

**Authors:** Heera Yoon, Myeong Seong Bak, Seung Ha Kim, Ji Hwan Lee, Geehoon Chung, Sang Jeong Kim, Sun Kwang Kim

## Abstract

We developed AI-bRNN (Average training, Individual test-bidirectional Recurrent Neural Network) to decipher spontaneous pain information from brain cellular calcium signals recorded by two-photon imaging in awake, head-fixed mice. The AI-bRNN determines the intensity and time point of spontaneous pain even during the chronic pain period and evaluates the efficacy of analgesics. Furthermore, it could be applied to different cell types and brain areas, and it distinguished between itch and pain, proving its versatility.

## Introduction

Spontaneous ongoing pain is a primary complaint in patients with chronic intractable pain^1^. However, the diagnosis and treatment of spontaneous pain remain challenging, partially because few objective methods are available to measure pain in preclinical research. In the last decade, inspiring approaches to spontaneous pain assessment have been adopted in animals. For example, the grimace scale (GS) detects facial expression to assess emotional response to pain^2^, while conditioned place preference (CPP) uses a motivational response to pain relief^3^. Although these methods have been successfully applied in several pain models, they have inevitable limitations. The GS cannot predict subchronic or chronic pain^2^, and CPP has limitations related to time resolution and cannot be used alongside drugs that impact reward response or learning and memory^4^. Thus, there is an unmet need for a new methodology to objectively evaluate spontaneous pain.

The neural processing of pain information involves multiple brain areas. Among these, the primary somatosensory (S1) cortex plays a key role in the perception and discrimination of pain sensation by encoding its intensity, location, and temporal course^5-7^. This led us to hypothesize that neuronal Ca^2+^ activity patterns in the mouse S1 cortex differ between spontaneous pain and non-pain conditions, and that spontaneous pain could be measured quantitatively based on this discrepancy.

## Results

To explore this hypothesis, we recorded S1 neuronal Ca^2+^ activity during formalin-induced spontaneous pain in awake, head-fixed mice (see Methods for details). Consistent with previous reports^8^, pain behaviors (i.e. licking and biting) were most prominent at 0-5 minutes after a formalin injection (5%, 10 μl, s.c.) into the right hind paw in freely moving mice (**Fig. 1a**, top right). Using *in vivo* two-photon microscopy and the genetically encoded Ca^2+^ indicator GCaMP6s, we imaged the Ca^2+^ activity of layer II/III neurons in the left S1 cortex of head-fixed mice while tracking their motion using an infrared camera (**Fig. 1a**, left). Only the Ca^2+^ signals recorded 1-3 minutes after the formalin injection were used as ‘pain’ condition signals (**Fig. 1a**, bottom right). We extracted raw Ca^2+^ traces from each region-of-interest (ROI) (**Fig. 1b**) and matched the heatmap-visualized Ca^2+^ traces (top) and the averaged trace (bottom) with the motion tracking data (**Fig. 1c**).

**Fig 1.**
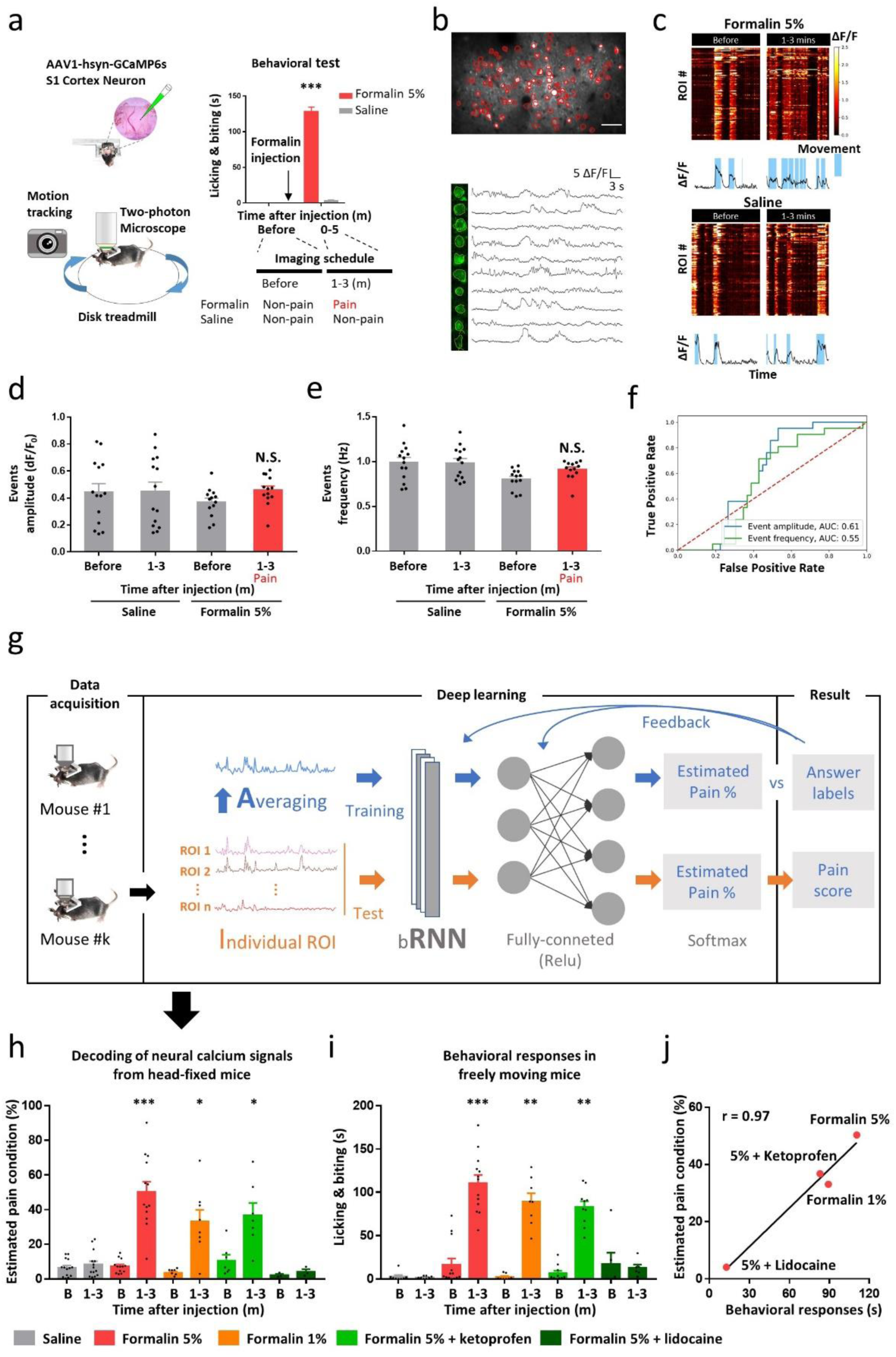
Conceptual framework of AI-bRNN and its validation. **a**, Schematic diagram of the experimental approach. AAV1-hsyn-GCaMP6s was injected into the S1 cortex (left). Imaging was performed for 2 minutes at each time point (before and 1-3 minutes after formalin injection). The time point of the imaging was determined based on the levels of nociceptive behavior after formalin injection in freely-moving animals (right). **b**, Representative image of S1 neurons identified by semi-automatic ROI analysis (top). Example of Ca^2+^ traces from each ROI (bottom). Scale bar represents 50 μm. **c**, Heatmaps showing the activity of S1 neurons. The line traces below each heatmap indicate the averaged values of the entire ROIs. The duration of the mouse movement identified by the motion tracking analysis was overlayed on the line trace using sky blue-shaded areas. **d, e**, Mean amplitude (**d**), and mean frequency (**e**) of Ca^2+^ events. **f**, Performance of the amplitude and frequency data for pain classification. **g**, Architecture of the AI-bRNN. The Ca^2+^ traces extracted from each ROI were averaged subject-by-subject to train the neural network. In the test session, the Ca^2+^ traces from individual ROIs were separately applied to the trained model. **h**, The prediction of the AI-bRNN regarding whether the subject was in a pain condition. On the x-axis, ‘B’ indicates before injection. Saline (s.c.) group (*n* = 14 mice); formalin 5% (s.c.) group (*n* = 13 mice); formalin 1% (s.c.) group (*n* = 8 mice); formalin 5% (s.c.) + ketoprofen (100 mg/kg, i.p.) group (*n* = 7 mice); formalin 5% (s.c.) + 2% lidocaine (10 μl, s.c.) group (*n* = 3 mice) **i**, Nociceptive behaviors in freely moving mice. Saline group (*n* = 8 mice); 5% formalin group (*n* = 13 mice); 1% formalin group (*n* = 8 mice); 5% formalin + ketoprofen group (*n* = 10 mice); 5% formalin+ 2% lidocaine group (*n* = 6 mice) **j**, Pearson’s correlation between the group average of the estimated pain values and the behavioral response time (Pearson’s r = 0.97). Scatter plots indicate individual data. Bars indicate mean ± s.e.m; N.S., non-significant; *** *P* < 0.001, ** *P* < 0.01, * *P* < 0.05 compared to baseline.

However, using conventional Ca^2+^ analyses^9^, we could not elicit pain-specific information from S1 neurons. The amplitude and frequency of Ca^2+^ events showed no differences between the pain and non-pain conditions (**Fig. 1d, e**), and the receiver-operating-characteristic (ROC) curve showed poor AUC scores of 0.61 and 0.55, respectively (**Fig. 1f**). These negative results may not be surprising, considering that S1 neurons process touch and proprioception as well as pain^7, 10^. Hence, we developed a deep learning algorithm to detect distinct features of Ca^2+^ activity that represent spontaneous pain.

We used a bidirectional recurrent neural network (bRNN), specialized for handling sequential data^11^, to identify spontaneous pain representation from thousands of sequential Ca^2+^ traces of S1 neurons. Through trial and error, we combined a preprocessing step that averages the Ca^2+^ traces from individual ROIs for training and leaves the individual traces for test with bRNN (AI-bRNN, **Fig. 1g**) to enhance the classification performance for various pain models (see **Extended Fig. 1**). Since the formalin pain model exhibits clear, strong, and measurable pain behaviors^12^ (**Fig. 1a**, top right), we utilized S1 neuronal Ca^2+^ signals recorded at 1-3 minutes following 5% formalin injection as supervisory signals for the AI-bRNN. After training the AI-bRNN, we tested its classification performance with the leave-one-subject-out (LOSO) cross-validation (CV) method. The AI-bRNN successfully predicted spontaneous pain conditions depending on the formalin concentration (5% and 1%; **Fig. 1h** and **Extended Fig. 1a**). Mild (ketoprofen, 100 mg/kg, i.p.) and strong (2% lidocaine, s.c.) painkillers reduced the estimated pain values in the 5% formalin group to the levels of the 1% formalin group and a saline-treated non-pain group, respectively (**Fig. 1h**). All of these pain values estimated by the AI-bRNN in head-fixed mice were similar to the pain behaviors measured in freely moving mice (**Fig. 1i**), with a highly positive correlation between the two (r = 0.97; **Fig. 1j**). These results indicate that our deep learning model can quantify the intensity of spontaneous pain and evaluate the efficacy of analgesics.

Next, we tested whether the AI-bRNN trained by S1 neuronal Ca^2+^ during formalin pain is broadly applicable to various pain models with different chronicities. Our deep learning algorithm discriminated another acute spontaneous pain induced by capsaicin (0.1%, s.c.) from non-pain condition (**Fig. 2a**). The AI-bRNN also distinguished subchronic spontaneous pain on days 1 and 3 after injection of complete Freund’s adjuvant (CFA; 10 μl, s.c.) (**Fig. 2a**). It is known that such pain cannot be indentified using GS^13^. Our motion tracking also failed to detect it (**Extended Fig. 2**). This indicates that the estimation of subchronic spontaneous pain by the AI-bRNN was independent of behavioral responses. Notably, the capsaicin and CFA pain data were completely isolated from the training session of the AI-bRNN, and these data were only used as test samples to avoid overfitting problems.

We further trained the AI-bRNN using additional Ca^2+^ data from the capsaicin/CFA group, but found that the resulting algorithm detected chronic neuropathic pain to a similar level as the algorithm trained using only the formalin group data (**Extended Fig. 1e**). This might be due to the less prominent pain-representative features in the capsaicin/CFA data. To circumvent this, we upgraded our deep learning model by implementing a semi-supervised learning strategy (**Fig. 2b**). Based on the formalin-induced Ca^2+^ signals, we assessed unlabeled Ca^2+^ data from the capsaicin and CFA groups to transform them into pseudo-labeled data. We then combined them with formalin-induced Ca^2+^ data for model training. This upgraded model clearly identified spontaneous neuropathic pain in the early (3d) and late (10d) phases following partial sciatic nerve ligation (PSL) injury (**Fig. 2c**), showing advanced classification performance (**Extended Fig. 1e**). Furthermore, we used sliding window method (segmentation of estimated pain values by 2-minute time windows) to reveal when, how long, and how much PSL group mice feel spontaneous pain (**Fig. 2d**). Combined treatment using gabapentin (GB; 100 mg/kg, i.p.) and venlafaxine (VX; 50 mg/kg, i.p.)^14^, which are clinically utilized analgesics, attenuated this spontaneous neuropathic pain at 3d, but not at 10d, following PSL (**Fig. 2c**,**d**), whereas it relieved stimulus-evoked pain (mechanical allodynia) in both phases (**Extended Fig. 3**). These results suggest that the AI-bRNN can be upgraded to provide more detailed information about spontaneous neuropathic pain, and that painkillers should be assessed separately for different pain types (i.e., evoked vs. spontaneous).

**Fig 2.**
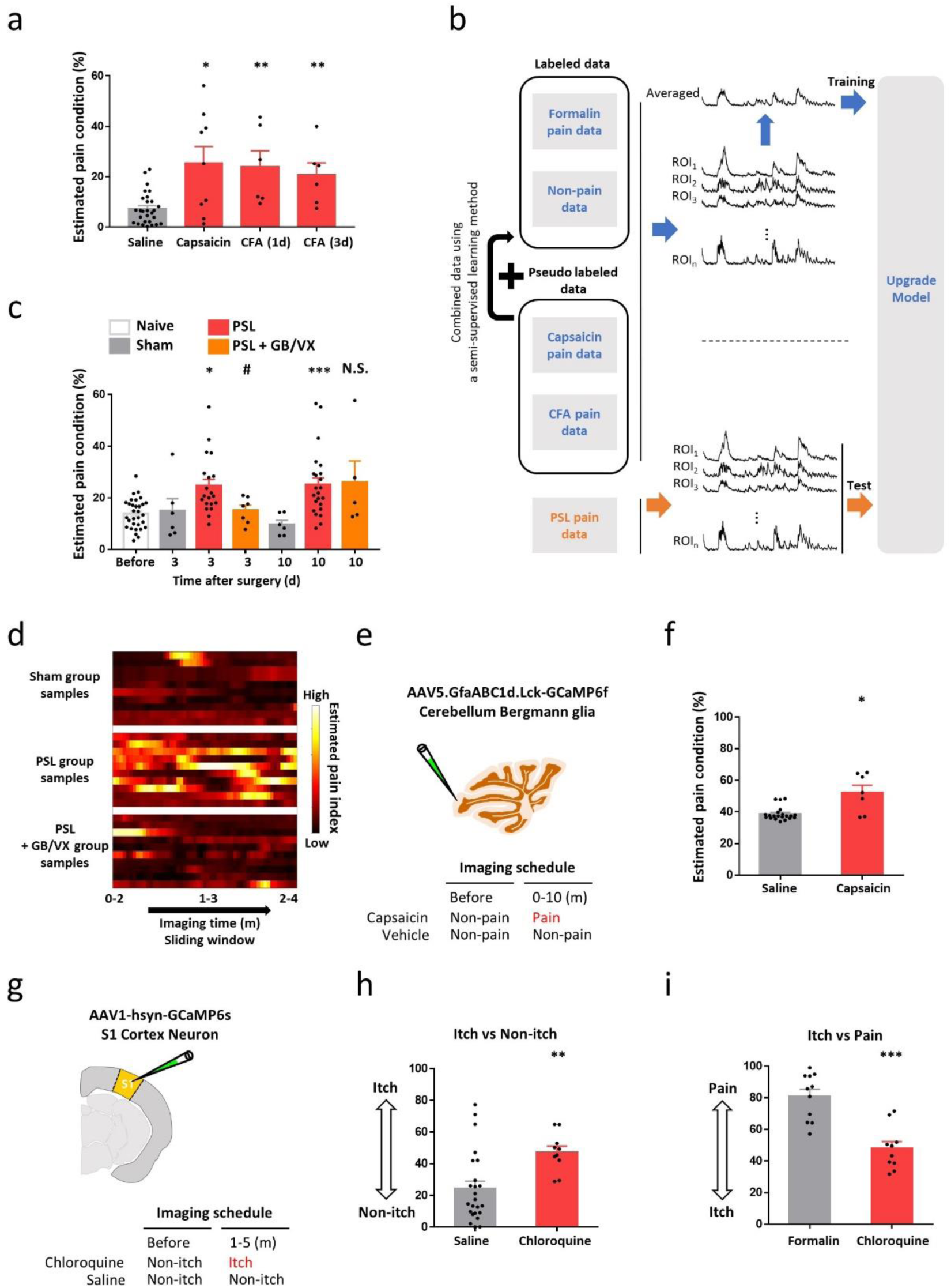
Broad applicability and versatile performance of AI-bRNN. **a**, Estimated pain values of the capsaicin or CFA-injected animals. The estimation was based on the Ca^2+^ activity of the neurons in the S1 cortex. Saline (s.c.) group (*n* = 28 sessions from 14 mice); capsaicin (0.01%, 10 μl, s.c.) group (*n* = 9 mice); CFA (10 μl, s.c.) group (*n* = 6 mice) **b**. Schematic diagram of semi-supervised learning method. Pseudo-labeled data were corrected based on formalin pain data. The upgraded model was trained with both labeled and pseudo-labeled data. PSL pain data were only used for the test session. **c**, Estimated pain values of the animals subjected to PSL or sham surgery. The estimation was based on the Ca^2+^ activity of the neurons in the S1 cortex. Sham group (*n* = 6 mice); PSL group (*n* = 20 mice); PSL + GB/VX (GB 100 mg/kg, VX 50 mg/kg, i.p.) group (*n* = 7 mice) **d**, Heatmap plot showing changes in estimated pain values over time with 2 minutes time resolution. **e**, Schematic drawing of cerebellum Bergmann glia and imaging timeline before and after capsaicin or vehicle injection. **f**, Estimated pain values of capsaicin-injected animals. The estimation was based on the Ca^2+^ activity of Bergmann glial cells in the cerebellum of the capsaicin group (*n* = 7 mice) and saline group (*n* = 14 mice). The data from baseline non-pain condition (before the capsaicin injection) were pooled with the data from the saline-injected animals. **g**, Schematic drawing of the Ca^2+^ imaging schedule of S1 neurons in chloroquine-induced itch conditions. **h**,**i**, Estimated itch values of the chloroquine-injected animals based on the Ca^2+^ activity of S1 neurons. The data from the baseline non-itch condition (before chloroquine injection) were pooled with the data from the saline-injected animals. Saline (10 μl, s.c.) group (*n* = 14 mice); chloroquine (100 μg/10 μl, s.c.) group (*n* = 10 mice); formalin (5%, s.c.) group (*n* = 11 mice). Scatter plots indicate individual data. Bars indicate mean ± s.e.m; *** *P* < 0.001, ** *P* < 0.01, * *P* < 0.05 compared to the matched control group, N.S., non-significant; ^#^ *P* < 0.05 compared to the matched non-pain group (Mann-Whitney test).

We then tested the versatile performance of the AI-bRNN by applying it to a different brain region (cerebellum) and cell type (glia), as well as to a different somatosensation (itch). Capsaicin-induced spontaneous pain information is transmitted to the cerebellum^15, 16^, leading to increased neuronal activity and subsequent Ca^2+^ elevation in the resident Bergmann glia (unpublished data). We expressed GCaMP6f in Bergmann glia of the cerebellar cortex lobule IV/V to image Ca^2+^ signals during capsaicin pain (0-10 minutes after injection) or non-pain conditions (**Fig. 2e**, see Methods for details). Since a region of Bergmann glia is hardly defined, we modified our deep learning model to use average Ca^2+^ signals from the whole imaging field, rather than individual Ca^2+^ traces from ROIs, in the training and test sessions. Nevertheless, the algorithm robustly estimated the capsaicin pain condition (**Fig. 2f, Extended Fig. 1d**). We also applied the AI-bRNN to classify another sensation, itch, which shares common neuroanatomical pathways with pain while being a clearly distinct sensation^17^. We acquired S1 neuronal Ca^2+^ signals during chloroquine (100 μl, s.c.)-induced spontaneous itch (1-5 minutes after injection) or non-itch conditions (**Fig. 2g**). The AI-bRNN successfully distinguished not only between itch and non-itch conditions, but also between pain and itch (**Fig. 2h, i** and **Extended Fig. 1d**).

## Discussion

Finding a new method that objectively and quantitatively measures spontaneous pain in animals is instrumental in unraveling the complicated neurobiological mechanisms of pain and discovering clinically relevant painkillers^18^. In the present study, we demonstrated that the AI-bRNN decoded detailed information about spontaneous pain, such as its intensity, time point and duration, based on brain cellular Ca^2+^ activity in awake mice. This cannot be achieved using existing methods. This deep learning algorithm is broadly applicable for assessing a wide range of spontaneous pain conditions (acute to chronic) and it could be used to validate the efficacy of analgesics. It should be noted that the novel analgesics have been discovered mostly by evoked pain tests in rodents, which are hardly translated into pain relief in human patients^19^. Thus, our unprecedented approach may offer a powerful preclinical evaluation platform for pain medicine.

Our approach also provides deep insight into the practical application of artificial intelligence in biology. Deep learning algorithms generally require massive, well-labeled data sets; biological data cannot usually meet this requirement^20^. To resolve this issue, we implemented several strategies as follows: First, we selected the formalin pain model to obtain supervisory Ca^2+^ signals as pain labels because this model produces robust and quantifiable spontaneous pain behaviors^8^; Secondly, we designed the deep learning algorithm, through trial and error, to use the averaged Ca^2+^ traces for training, which enhances its classification performance. Thirdly, our algorithm could be flexibly modified depending on the property and shape of the available Ca^2+^ data. We upgraded the AI-bRNN to detect chronic neuropathic pain by combining a semi-supervised learning paradigm, which provides pseudo-labeled data from an unlabeled data set and eventually improves performance. We also modified the algorithm (AI-bRNN→AA-bRNN) to apply it in the cerebellar Bergmann glia, which could not be separated by individual ROIs. Additionally, we demonstrated that the AI-bRNN can be applied to other types of sensation, such as itch, further distinguishing between itch and pain. Such flexibility, broad applicability and versatile performance are partially underpinned by the RNN’s competence, which automatically extracts features from sequential data such as Ca^2+^ imaging data, irrespective of prior knowledge from a specific domain^11^. Taken together, we propose that our approach may help decode not only brain cellular Ca^2+^ signals, but also other forms of sequential data in biology.

## Author contributions

SJK and SKK conceived and supervised the project. HY, MSB, GC, SJK and SKK designed the study. HY performed all the two-photon imaging experiments in the S1 cortex in awake head-fixed mice and the behavioral tests in freely moving mice. MSB developed the deep learning algorithm (AI-bRNN) with intellectual inputs from GC and SKK, and processed and analyzed all the Ca^2+^ imaging data. SHK conducted the two-photon imaging experiments on Bergmann glia in the cerebellum. JHL performed and analyzed all the behavioral tests in freely moving mice. HY, MSB, GC and SKK wrote the manuscript with comments from SJK. All authors read and approved the manuscript.

## Acknowledgements

We thank Changmin Oh and Yangseok Kim for their valuable advice on deep learning algorithms; Yoorim Kim, Yong-Seok Lee, Hyunsu Bae and Sun Wook Hwang for critical discussions on the experiments; Junichi Nabekura and Schuichi Koizumi for providing cover glasses and encouragement. This work was supported by grants from the National Research Foundation of Korea funded by the Korean government (NRF-2017M3C7A1025604, NRF-2017M3A9E4057926, and NRF-2019R1A2C2086052 to SKK; NRF-2018R1A5A2025964 and NRF-2017M3C7A1029611 to SJK).

## Online Methods

### Experimental animals

All mice were C57BL/6 male, aged 5-6 weeks old at the start of the experiments. The mice were housed in groups of two to minimize stress. The vivarium was controlled with a 12-h light/dark cycle, and all experiments were performed during the daylight hours. All experimental procedures were approved by the Seoul National University Institutional Animal Care and Use Committee and performed in accordance with the guidelines of the National Institutes of Health.

### Behavior test

The formalin test was performed in mice that had been individually exposed to the observation chamber. Before the test, the mice were acclimated to the chamber for 1 h on 3 consecutive days. Ten μl of 5% or 1% formalin solution was injected into the right hind paw; the mouse was then immediately put back into the chamber. We recorded the time spent in nociceptive behavior (licking and biting of the injected paw) in each 5 minutes-period. In the experiments involving analgesic drugs, ketoprofen (100 mg/kg, 50 μl, i.p.) or 2% lidocaine (10 μl, subplantar, s.c.) was administered 20 minutes before the formalin injection.

To quantify the relieving effect of gabapentin/venlafaxine (GB/VX) on PSL-induced neuropathic mechanical allodynia, the mice were placed in a 12 (d) × 8 (w) × 6 (h) cm clear plastic cage on a metal mesh. Behavioral tests were performed 30, 60, and 120 minutes after GB (100 mg/kg) and VX (50 mg/kg) administration. Mechanical allodynia was assessed using the von Frey filament (Linton Instrumentation, Norfolk, UK). Specifically, von Frey filaments delivering a different bending force (2.36, 2.44, 2.83, 3.22, 3.61, 3.84, 4.08, and 4.31) were applied to the right hind paw using the up-down method, and the threshold force corresponding to 50% withdrawal was determined^21^. The experimenters were blinded to the treatment information of the animals.

### Surgical preparation for imaging awake mice

All surgical procedures were performed under isoflurane anesthesia (1-1.5%). To minimize edema and related inflammation, dexamethasone (0.2 mg/kg) and meloxicam (20 mg/kg) were administered by subcutaneous injection. A cranial window was made above the left S1 cortical hind paw area (size: 2 × 2 mm: center relative to Bregma: lateral, 1.5 mm, posterior, 0.5 mm)^22, 23^. A small craniotomy was carefully performed using a #11 surgical blade. The exposed cortex was perfused with artificial cerebrospinal fluid, and adeno-associated virus expressing GCaMP6s (AV-1-PV2824; produced by the University of Pennsylvania Gene Therapy Program Vector Core) was injected into the S1 cortex (30-50 nl per site; 200-300 μm from the surface) using a broken glass electrode (20-40 μm tip diameter). After virus injection, the exposed cortex was covered with a thin cover glass (Matsunami, Japan) and the margin between the skull and the cover glass was tightly sealed using Vetbond (3M) and dental cement. For cerebellar Bergmann glia imaging, a small craniotomy was performed over the lobule IV/V cerebellar vermis. AAV5.GfaABC1d.Lck-GCaMP6f was delivered to the cerebellum (100-300 μm from the surface), as described above.

### Two-photon Ca^2+^ imaging of awake mice

For awake imaging, mice were habituated on the treadmill under head-fixed conditions for 40 minutes per day over 2 weeks. Ca^2+^ imaging was performed using a two-photon microscope (Zeiss LSM 7 MP, Carl Zeiss, Jena, Germany) equipped with a water immersion objective lens (Apochromat 20, NA = 1.0, Carl Zeiss). Two-photon excitation at 900nm for GCaMP6s imaging was carried out using a mode-locked Ti: sapphire laser system (Chameleon, Coherent, USA). Data were acquired using ZEN software (Zeiss Efficient Navigation, Carl Zeiss) at 4.4 Hz for S1 imaging and 32 Hz for cerebellar Bergmann glia imaging.

### Motion tracking during calcium imaging

Mouse locomotion was recorded using a video camera. Motion tracking was controlled by a custom program written in LabVIEW (National Instruments, USA) and was synchronized with two-photon imaging by a trigger generated in the program. Mouse locomotion was recorded at a rate of 64 Hz using a high-speed CCD camera (IPX-VGA210, IMPERX, USA) with infrared illumination (DR4-56R-IR85, LVS, S. Korea).The recorded video had 64 frames per second, with a frame size of 720 x 480 pixels. To assess the level of locomotion, the difference in intensity of each pixel between frames was calculated and summarized across all the pixels. If the summarized value of a frame exceeded an arbitrary threshold determined by a experimenter blind to the treatment information, the frame was scored as a locomotion-positive frame.

### Experimental models of spontaneous pain

For the formalin-induced pain model, 10 μl of formalin (5% or 1%) or vehicle solution was injected into the right hind paw. For the capsaicin model, 10 μl of capsaicin (0.01%) was delivered to the right hind paw. For the CFA model, 10 μl of CFA was injected subcutaneously into the plantar surface of the right hind paw. For the neuropathic pain model, PSL surgery was performed to the right hind paw under isoflurane anesthesia. The right sciatic nerve was exposed at the upper thigh of the mouse, and one-third to one-half of the nerve diameter was ligated with a 9-0 suture. In the assessment of the analgesic effects, GB (100 mg/kg) and VX (50 mg/kg) were co-administered to the PSL animals 3 and 10 days after the surgery. Imaging and behavioral tests were performed 30 minutes after the intraperitoneal injection of the drugs. For the itch model, chloroquine (100 μg/10 μl) was delivered to the right hind paw. Imaging was performed 1-5 minutes after chloroquine administration.

### Extracting Ca^2+^ traces and event detection

The imaging data were motion-corrected using the Turboreg algorithm (Biomedical Imaging Group, Swiss Federal Institute of Technology, Lausanne, Switzerland). The ROIs were detected using the CNMF-E algorithm^24^ and manually reviewed. The spatial information of the ROI was imported into ImageJ (https://imagej.nih.gov/ij/) and the average fluorescence of the ROI was calculated along with the frame. The events were detected from the extracted Ca^2+^ traces using MLSpike^9^, an open-source software. The hyperparameters were set as follows: a = 0.3, tau = 1, saturation = 0.1, finetune.sigma = 0.02, and drift.parameter = 0.1. The drift signal from MLSpike was used to calculate the amplitude and frequency of the events.

### Ca^2+^ signal normalization

A Gaussian window^25^ (10 frames in extent) was applied to smooth the extracted Ca^2+^ traces. The signals below the 70^th^ percentile in each ROI were averaged and used as a baseline fluorescence signal (F0). All signals were transformed to dF/F0 in each ROI to normalize the scale range^7^. The data length was fixed to 497 frames. If the data of a subject had more than 497 frames, we applied the window slicing method (window size of 497 frames, step size of 10 frames) and used the average of all window slides as the final score.

### Preprocessing for deep learning and deep learning architecture

The Ca^2+^ imaging data obtained in one imaging session had a size of n x m, where n was the number of ROIs and m was the number of frames (typically 497 frames for 2 minutes). The deep learning model was trained to have an output of [1, 0] for the non-pain condition or a [0, 1] for the pain condition. The neural network was implemented using Keras^26^, The source codes are available from https://github.com/KHUSKlab/ paindecoding. In step (i), the first input to the neural network had the shape (k, 497, 1), where k is the number of input data in the training set, 497 is the frame length of the time series Ca^2+^ signals used, and 1 is the number of features. In step (ii), this input was forwarded to the bidirectional recurrent neural network^11^ (long short-trem memory cells^27^) and activated by a hyperbolic tangent function. Next, in step (iii), the data were forwarded to two dense layers, with drop out rates of 0.2 and 0.1, respectively. These two dense layers were activated by rectified linear unit (RELU)^28^ and sigmoid functions, respectively. Finally, the data were forwarded to a dense layer that was activated by the softmax function, defined as follows;

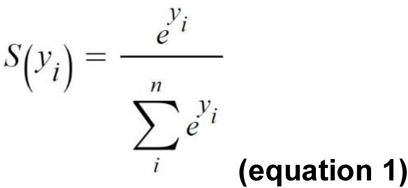

where y is the activated value, i is each node (i.e., class), and n is the total number of nodes or classes. The loss function was defined as follows;

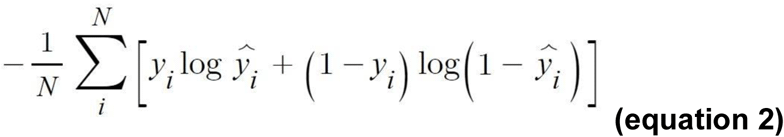

where y hat is estimated value, which was activated by the softmax function, y is the label [1,0] for non-pain or [0, 1] for pain conditions, and N is the total number of samples. The Adam optimizer^29^ was used and the hyperparameters were set as default (lr = 0.01, decay = 1e-8, β1 = 0.9, β2 = 0.999). The L2 regularizer was applied to layers in step (iii) to prevent overfitting. Therefore, the loss function was redefined as follows;

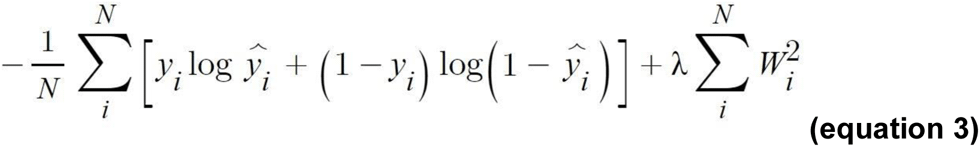

where W is the summation of the weight values of the layers in step (iii), and λ is the weight of the l2 regularizer. The initial weights of all layers were set by He uniform variance scaling initializer with a fixed random seed to ensure reproducibility.

### Training and test of the deep learning model

In this study, several models were used depending on the classification purpose and the shape of the data. Therefore, each model is described separately below. Generally, the number of pain (or itch) samples for deep learning training was less than the number of non-pain samples. To balance each class, the pain (or itch) samples were duplicated.

### The deep learning model for formalin, capsaicin, and CFA pain

To estimate formalin-induced pain or non-pain samples, we used leave-one-subject-out (LOSO) cross-validation (CV). Thus, all data obtained from one mouse (subject) were isolated and assigned only to the test session but not to the training session. In the case of other samples (capsaicin- and CFA-induced pain group), data were never used to train the deep learning model and were tested as out-of-sample.

### The deep learning model for PSL-induced pain

As described above, semi-supervised learning paradigms were adopted to upgrade the deep learning model. Ca^2+^ traces from capsaicin and CFA groups were sampled and randomly selected to create a subset for model training. The CFA group was subjected to 4 minutes of imaging (1,045 frames), compared to the 2 minutes (497 frames) in the other groups, so 55 subsamples were extracted from each CFA-injected subject using the window-slicing method. A temporary deep learning model was trained using a randomly chosen subset of the capsaicin and resampled CFA data, and data from the formalin group were then estimated using this temporary model. The total classification accuracy of the model for formalin pain data was evaluated. In bootstrapping, the contribution of a sample to the classification performance was evaluated by averaging the accuracy values of the subsets that included the sample. If a sample of the capsaicin and CFA group produced sufficient contribution to the classification accuracy (average accuracy of the subsets > 70%), it was chosen as a pseudo-labeled sample. All pseudo-labeled data were combined with formalin pain data to train the deep learning model in the next step and the PSL data were estimated using this upgraded model. The samples of the PSL pain group were never used for deep learning training, so PSL pain signals were tested as out-of-sample.

### The deep learning model using Bergmann glia signal

The Ca^2+^ imaging data of the cerebellum Bergmann glia (19200 frames per subject) were downsampled to 1/40 (479 frames). Additionally, the deep learning model received mouse locomotion data in parallel, so the input to the neural network had the shape (k, 479, 2). For the test, the LOSO-CV method was applied.

### The deep learning model for chloroquine-induced itch

To discriminate itch and non-itch signals, we trained a deep learning model using Ca^2+^ imaging data obtained from chloroquine-injected animals and saline-injected animals. To discriminate itch (non-pain) and pain signals, we reused the established deep learning model that had been trained using data from the formalin-injected animals. Samples used in the training session were tested with the LOSO-CV method. Samples not used in the training session were tested as out-of-sample.

### Statistics

Statistical analyses were performed using GraphPad Prism 7 (GraphPad Software, Inc.) or Python (scipy library^30^). Two-factor repeated-measure ANOVA and Student-Neuman-Keuls *post hoc* were used in Extended Fig. 3. Pearson correlation analysis was used in Fig. 1j. Wilcoxon test was used in Fig. 1h. Mann-Whitney test was used in all other analyses. All values are represented as mean ± s.e.m.

---

**Extended Fig. 1.**
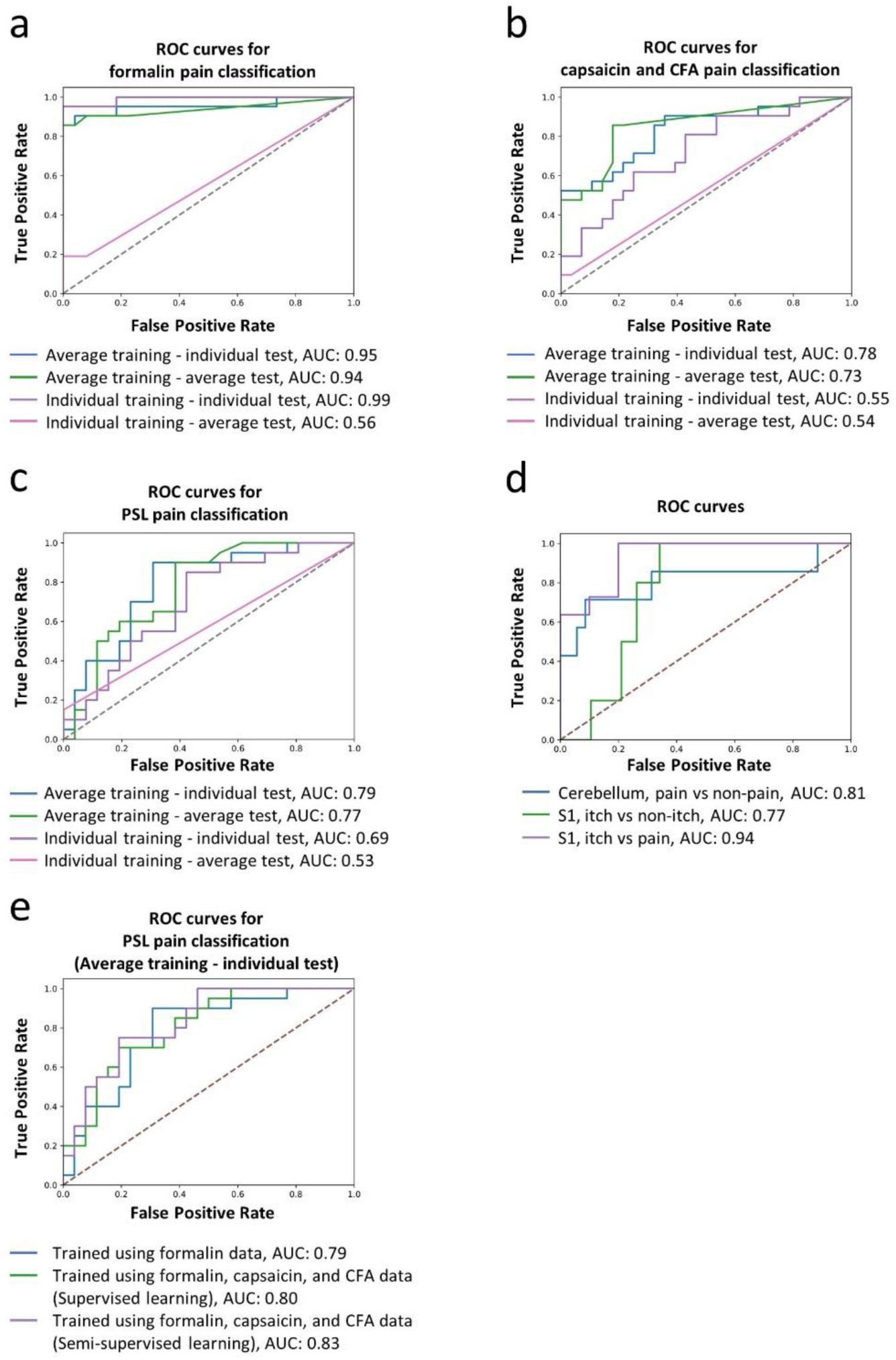
Classification performance of deep learning models with different pre-processing. Comparison of the area under the ROC curves for the four types of training-test strategies in each pain model. **a-c**, Classification performance for formalin (**a**), capsaicin, CFA (**b**) or PSL (**c**) pain conditions based on the S1 cortex neural signals. **d**. Classification performance for capsaicin-induced pain conditions based on the cerebellar Bergmann glia signals and for chloroquine-induced itch conditions based on the S1 cortex neural signals.

**Extended Fig. 2.**
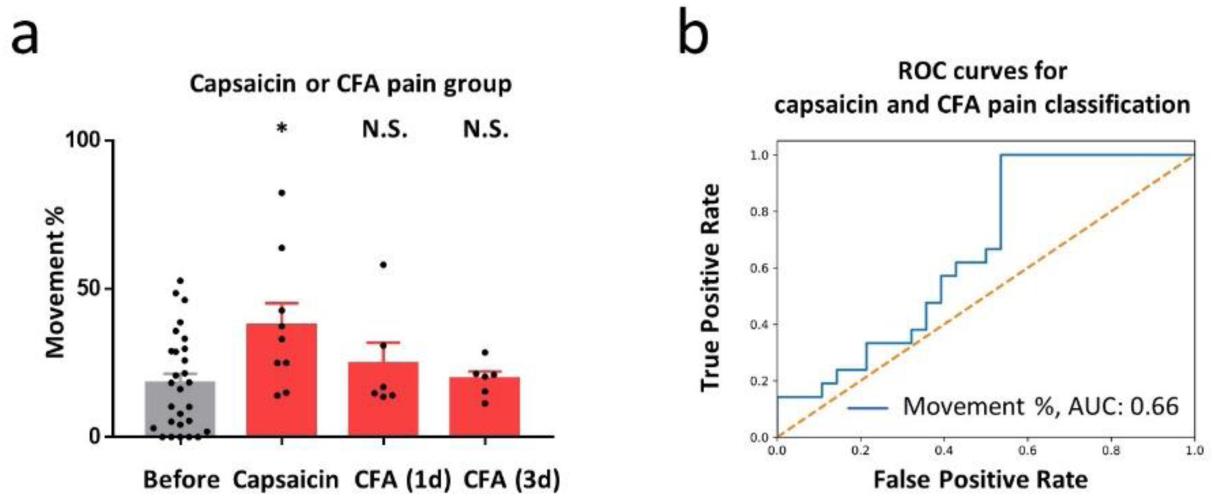
Locomotion of awake, head-fixed mice recorded by motion tracking analysis. **a**, Locomotion of the capsaicin- or CFA-injected animals. Bars indicate mean ± s.e.m. Scatter plots indicate individual data. **b**, Classification performance for capsaicin- or CFA-induced pain conditions based on locomotion levels. N.S., non-significant; ^*^ *P* < 0.05 compared to the matched non-pain group (Mann-Whitney test).

**Extended Fig. 3.**
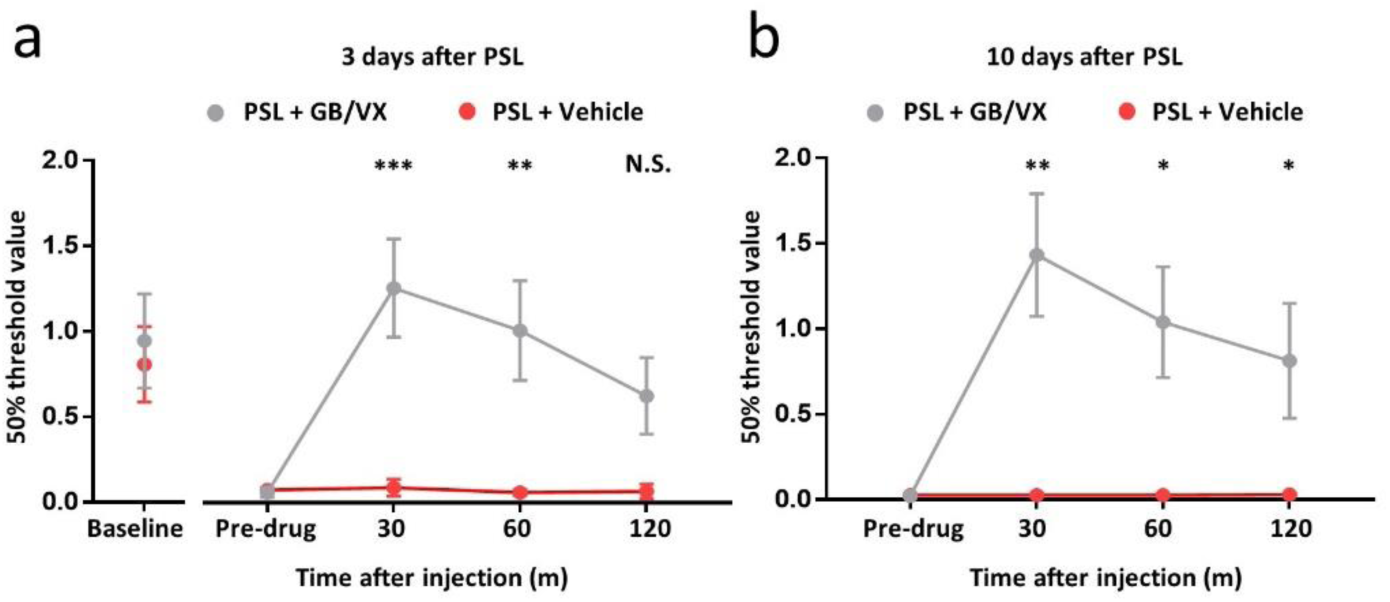
Relieving effects of GB/VX on neuropathic mechanical allodynia. Paw withdrawal threshold after GB and VX treatment on (**a**) days 3 and (**b**) 10 after PSL surgery in freely moving mice. Plots indicate mean ± s.e.m; *** *P* < 0.001, ** *P* < 0.01, * *P* < 0.05 compared to pre-drug values (two-factor repeated-measure ANOVA, Student-Neuman-Keuls *post hoc* test).

## References

1. Backonja, M.M. & Stacey, B. Neuropathic pain symptoms relative to overall pain rating. J Pain 5, 491–497 (2004).

2. Langford, D.J. et al. Coding of facial expressions of pain in the laboratory mouse. Nat Methods 7, 447–449 (2010).

3. King, T. et al. Unmasking the tonic-aversive state in neuropathic pain. Nat Neurosci 12, 1364–1366 (2009).

4. Tappe-Theodor, A., King, T. & Morgan, M.M. Pros and Cons of Clinically Relevant Methods to Assess Pain in Rodents. Neurosci Biobehav Rev 100, 335–343 (2019).

5. Bushnell, M.C. et al. Pain perception: is there a role for primary somatosensory cortex? Proc Natl Acad Sci U S A 96, 7705–7709 (1999).

6. Rocchi, L., Casula, E., Tocco, P., Berardelli, A. & Rothwell, J. Somatosensory Temporal Discrimination Threshold Involves Inhibitory Mechanisms in the Primary Somatosensory Area. J Neurosci 36, 325–335 (2016).

7. Kim, Y.R., Kim, C.E., Yoon, H., Kim, S.K. & Kim, S.J. S1 Employs Feature-Dependent Differential Selectivity of Single Cells and Distributed Patterns of Populations to Encode Mechanosensations. Front Cell Neurosci 13, 132 (2019).

8. Dubuisson, D. & Dennis, S.G. The formalin test: a quantitative study of the analgesic effects of morphine, meperidine, and brain stem stimulation in rats and cats. Pain 4, 161–174 (1977).

9. Deneux, T. et al. Accurate spike estimation from noisy calcium signals for ultrafast three-dimensional imaging of large neuronal populations in vivo. Nature Communications 7, 12190 (2016).

10. Kim, S.S., Gomez-Ramirez, M., Thakur, P.H. & Hsiao, S.S. Multimodal Interactions between Proprioceptive and Cutaneous Signals in Primary Somatosensory Cortex. Neuron 86, 555–566 (2015).

11. Schuster, M. & Paliwal, K.K. Bidirectional recurrent neural networks. IEEE Trans. Sig. Proc. 45, 2673–2681 (1997).

12. Hunskaar, S., Fasmer, O.B. & Hole, K. Formalin test in mice, a useful technique for evaluating mild analgesics. J Neurosci Methods 14, 69–76 (1985).

13. Sotocinal, S.G. et al. The Rat Grimace Scale: a partially automated method for quantifying pain in the laboratory rat via facial expressions. Mol Pain 7, 55 (2011).

14. Rode, F., Brolos, T., Blackburn-Munro, G. & Bjerrum, O.J. Venlafaxine compromises the antinociceptive actions of gabapentin in rat models of neuropathic and persistent pain. Psychopharmacology 187, 364–375 (2006).

15. Iadarola, M.J. et al. Neural activation during acute capsaicin-evoked pain and allodynia assessed with PET. Brain 121 931–947 (1998).

16. Moulton, E.A., Schmahmann, J.D., Becerra, L. & Borsook, D. The cerebellum and pain: passive integrator or active participator? Brain Res Rev 65, 14–27 (2010).

17. Dhand, A. & Aminoff, M.J. The neurology of itch. Brain 137, 313–322 (2014).

18. Xu, B., Descalzi, G., Ye, H.-R., Zhuo, M. & Wang, Y.-W. Translational investigation and treatment of neuropathic pain. Molecular Pain 8, 15 (2012).

19. Vierck, C.J., Hansson, P.T. & Yezierski, R.P. Clinical and pre-clinical pain assessment: are we measuring the same thing? Pain 135, 7–10 (2008).

20. Webb, S. Deep learning for biology. Nature 554, 555–557 (2018).

21. Chaplan, S.R., Bach, F.W., Pogrel, J.W., Chung, J.M. & Yaksh, T.L. Quantitative assessment of tactile allodynia in the rat paw. J Neurosci Methods 53, 55–63 (1994).

22. Kim, S.K. & Nabekura, J. Rapid synaptic remodeling in the adult somatosensory cortex following peripheral nerve injury and its association with neuropathic pain. J Neurosci 31, 5477–5482 (2011).

23. Eto, K. et al. Inter-regional contribution of enhanced activity of the primary somatosensory cortex to the anterior cingulate cortex accelerates chronic pain behavior. J Neurosci 31, 7631–7636 (2011).

24. Zhou, P. et al. Efficient and accurate extraction of in vivo calcium signals from microendoscopic video data. Elife 7, e28728 (2018).

25. Oppenheim, S. Schafer, Ronald W. Discrete-Time Signal Processing. 468–471 (1999).

26. Chollet, F. Keras. https://github.com/fchollet/keras (2015).

27. Hochreiter, S. & Schmidhuber, J. Long short-term memory. Neural Computation 9, 1735–1780 (1997).

28. Glorot, X., Bordes, A. & Bengio, Y. in Proceedings of the Fourteenth International Conference on Artificial Intelligence and Statistics, Vol. 15. (eds. G. Geoffrey, D. David & D. Miroslav) 315-323 (PMLR, Proceedings of Machine Learning Research; 2011).

29. Kingma, D. & Ba, J. Adam: A Method for Stochastic Optimization. International Conference on Learning Representations (2014).

30. Jones, E., Oliphant, T and Peterson, P ‘SciPy: Open source scientific tools for Python,’ URL: http://www.scipy.org/ (2001).

